# BI-SCUDO Regression - A novel algorithm to infer clinical variables from omics datasets

**DOI:** 10.64898/2026.11.17.767338

**Authors:** Giacomo Fantoni, Mario Lauria, Carlo Alberto Rossi, Roberto Bizzotto, Luca Marchetti

## Abstract

Regression analysis of high-throughput omics data is often made difficult by noise, batch effects, and the loss of information resulting from the discretization of continuous variables. Existing machine learning algorithms often struggle to extract stable biological signals under these heterogeneous conditions.

We introduce BI-SCUDO Regression, a rank-based regression framework designed for omics data analysis. By leveraging within-sample rankings and network-based predictions, BI-SCUDO Regression bypasses the need for complex normalization and provides robust, interpretable results. In synthetic benchmarks, BI-SCUDO Regression demonstrated strong resilience to noise, maintaining predictive and feature-selection stability under conditions in which standard models exhibited performance degradation. When applied to cohorts from TCGA datasets, BI-SCUDO Regression consistently outperformed LASSO, Random Forest Regression, and XGBoost in predicting survival, with an average increase in Ratio of Performance to Deviation ranging from 43% to 86%, depending on the comparator. Furthermore, the algorithm consistently identified parsimonious, statistically significant, and biologically relevant biomarkers, with enrichment for cancer-related terms across all stage stratifications. Although BI-SCUDO Regression incurs higher training times due to its genetic optimization strategy, it offers a robust and interpretable approach to outcome prediction and biomarker discovery, with potential applications in precision medicine, where noisy data compromise the performance of conventional regression methods.

## Introduction

Next-generation sequencing and other high-throughput technologies have transformed biomedical research by enabling the rapid generation of large and heterogeneous omics datasets [1]. Analyses of these data have already driven advances in disease diagnosis, prognosis, and personalized medicine. Yet, deriving insights from such datasets remains a central challenge in bioinformatics. Due to their highly dimensional, noisy, and heterogeneous nature, extracting reliable, clinically meaningful predictions is a complex task [2].

Classical machine learning methods such as LASSO [3], Random Forest Regression [4], and XGBoost [5] have been widely applied to omics regression tasks with variable results [6]. While attractive for their relative simplicity, these approaches often struggle with overfitting, multicollinearity, and the curse of dimensionality [7] when applied to datasets with thousands of features and a limited sample size. As a result, predictive accuracy and stability are often compromised when targeting continuous clinical outcomes.

A common workaround is to discretize continuous phenotypes into categorical labels, converting regression problems into classification tasks [8]. This reduction simplifies model training, reduces computational complexity, and can yield improved performance. However, it comes at the cost of discarding the fine-grained information found in continuous outcomes. Such information is essential for precision medicine, which aims to provide patient-specific predictions rather than coarse groupings, limiting the applicability of this strategy in these settings.

Deep learning methods have also been explored for omics regression, often achieving better predictive performance than classical algorithms [9]. However, the complexity of these architectures makes them difficult to interpret, rendering them poorly suited for clinical and translational research, where interpretability is not optional: physicians and researchers require not only accurate predictions but also an understanding of the molecular features driving them. Although post-hoc explainability methods exist to interpret complex deep learning models, their explanations can be approximate and less reliable [10]. Therefore, the limited transparency of deep learning models constrains their practical adoption in precision medicine.

Another recurring challenge for both classical and modern methods is robustness to dataset heterogeneity. Omics data collected across laboratories, platforms, or patient cohorts are often affected by batch effects [11]. These technical variations can overshadow biological signals, leading to poor generalization when models are applied to new samples or independent datasets. While normalization procedures are routinely employed, they introduce additional preprocessing steps and are not always effective, particularly when the distributions of external data differ substantially from those of the training data.

These limitations underscore the need for regression methods that can directly address the complexities of high-dimensional omics data without sacrificing interpretability or robustness. To this end, we propose BI-SCUDO Regression (Biomarker Identification – Signature based ClUstering for Diagnostic purpOses), a novel rank-based framework designed for continuous outcome prediction in heterogeneous omics and clinical datasets. While built upon the mathematically sound foundation of the SCUDO [12] classification algorithm, a methodology extensively validated across a broad and diverse omics applications ([13], [14], [15], [16], [17], [18]), BI-SCUDO

Regression is a distinct paradigm shift specifically designed for continuous outcomes. The innovation of this method rests on three key innovations: it employs rank transformation [19] to stabilize variance and reduce sensitivity to noise while preserving the granularity of continuous outcomes, it integrates robust feature selection to mitigate the effects of high dimensionality and highlight the most informative signals, and, finally, its network-based design provides intuitive visualization of model outputs and enhances interpretability.

A distinctive feature of BI-SCUDO Regression is its ability to extract discriminative features on a per-sample basis. Instead of generating population-level associations alone, the method identifies molecular characteristics directly linked to the clinical variable for each individual. This property is particularly important for translational applications, where clinicians need to understand the drivers of predictions at the level of individual patients.

By combining predictive accuracy, interpretability, and robustness, BI-SCUDO Regression directly addresses critical shortcomings of existing approaches. In doing so, it provides a practical tool for analyzing high-dimensional omics data in precision medicine, enabling fine-grained, clinically relevant predictions that better support personalized care.

In this study, we introduce BI-SCUDO Regression and systematically evaluate its performance. We first validate the method on synthetic datasets to assess accuracy, robustness, and interpretability under controlled conditions. We then demonstrate its applicability to real-world settings by analyzing all the publicly available transcriptomic data from The Cancer Genome Atlas (TCGA) [20]. Together, these results highlight BI-SCUDO Regression as a practical and interpretable framework for continuous outcome prediction in heterogeneous omics datasets.

## Materials and Methods

In this section, we provide a technical description and mathematical formulation of how BI-SCUDO Regression builds the relationship between samples, how we generated and collected the data used for testing, and how we compared it with state-of-the-art methods. BI-SCUDO Regression comprises four major steps: (i) signature extraction, (ii) similarity graph construction, (iii) optimization, and (iv) prediction. While a comprehensive description of the full pipeline is provided in Section 3.1, this section details the mathematical formulation of the algorithm.

### Extracting Patient-Specific Signatures

BI-SCUDO Regression employs a multi-step procedure to derive robust, patient-specific signatures from high-dimensional omics data. Given a training cohort with a continuous dependent variable *Y* ∈ ℝ^*m*^, where *m* is the number of samples, firstly BI-SCUDO Regression identifies the “low” (*S*_*low*_) and “high” (*S*_*high*_) proxy classes based on the extreme *P*_*i*_-percentiles of the distribution of *Y* . Then a univariate Wilcoxon rank-sum test is applied to each feature to evaluate the differential expression of features between *S*_*low*_ and *S*_*high*_. A dynamic threshold *p*_*thresh*_ is defined so that *p*_*thresh*_ ∈ [0, 0.05]. Any feature whose p-value *p > p*_*thresh*_ are discarded to remove features lacking response-dependent variation, like housekeeping genes for transcriptomic data. The remaining features are rank transformed: absolute expression values are replaced with within-sample ranks, denoted as *r*_*f,s*_ for feature *f* in sample *s*. This nonparametric approach relies on relative rankings rather than precise expression values, preserving the within-sample ordering of gene expression while improving model robustness to inter-batch technical variation and cross-sample normalization artifacts. Finally, for each sample *s*, the *n*_1_ highest-ranked and *n*_2_ lowest-ranked features are extracted. The resulting patient-specific signature *R*_*s*_ is a discrete feature set of length *n*1 + *n*2. The signature represents a two-sided snapshot of the sample’s state and captures both active drivers and suppressed pathways that characterize the sample’s unique behavior.

### Building the BI-SCUDO Regression similarity graph

The similarity graph is the underlying data structure of BI-SCUDO Regression. Nodes *{*1, *…, m}* represent training samples, and edges encode pairwise similarities weighted by the inverse of a symmetric similarity metric *D* (defined below). Local connectivity is ensured by defining the directed adjacency matrix *a*_*i,j*_ as:

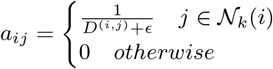

Where *i* and *j* are nodes, *ϵ* is a small smoothing constant and:

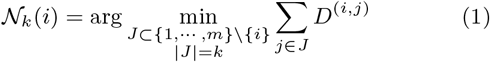

is the *k*-nearest neighbors set of sample *i*. To facilitate convex averaging during prediction, the weights outgoing from each node *i* are normalized to sum to one:

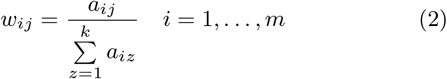

Finally, the distance *D*^(*i,j*)^ is computed via a weighted enrichment score [21] procedure applied to the feature signatures *R*_*i*_ and *R*_*j*_ obtained following the procedure in Section 2.1. The weight assigned to each feature reflects its discriminative power between the *S*_*low*_ and *S*_*high*_ classes, as defined in Section 2.1. Let *r*_*f,s*_ denote the rank of feature *f* in sample *s*. The median within-class rank for feature *f* in class *c* ∈ *{low, high}* is:

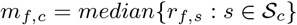

The within-class rank range is defined as:

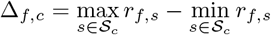

To quantify feature importance, we calculated the inter-class separation (*δ*_*f*_) as the absolute difference between class medians, and the intra-class variability (Δ_*f*_) as the average of the two within-class rank ranges.

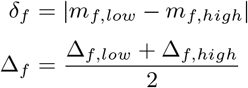

The unnormalized weight *u*_*f*_ is then expressed as the ratio of separation to variability:

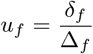

which is normalized between [0, 1] as 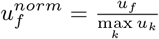 . This weighting scheme prioritizes features characterized by high inter-class divergence and consistent within-class behavior, enhancing the robustness of the enrichment score calculations.

### Prediction and Optimization

Once the similarity graph is built, BI-SCUDO Regression makes a prediction for a new, unseen sample *s*^*^ by first extracting its signature _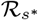_ and then exploiting it to compute the pairwise distance 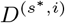 with all the samples *i* in the training set. Based on the distance values, the sample is added to the similarity graph by including an edge from it to its *k* nearest neighbors, identified by equation 1, with a weight 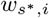, as computed by equation 2. The predicted dependent variable 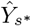 is predicted as:

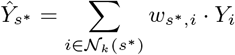

This nonparametric approach ensures that the prediction is determined by the neighborhood of patients who share the most similar molecular signature, providing high interpretability. To ensure that the model is robust, parsimonious, and makes the best prediction possible, BI-SCUDO Regression integrates a parameter optimization procedure based on a genetic algorithm to search the parameter space Θ = *{P*_*i*_, *p*_*thresh*_, *n*_1_, *n*_2_, *k}* to minimize:

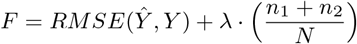

Where RMSE is the Root Mean Square Error of the prediction, the second term serves as a parsimony penalty for regularization, *N* is the total number of features, and *λ* is a scaling factor heuristically computed to allow the selection of parsimonious biomarkers while preserving the optimality of the predictions. This term forces the algorithm to favor smaller, highly specific gene signatures over large, noisy sets.

## Comparison with the state-of-the-art

To evaluate the performance and interpretability of BI-SCUDO Regression, we benchmarked it against three state-of-the-art explainable regression frameworks: LASSO [3], Random Forest Regression [4], and XGBoost [5]. These methods represent a diverse suite of algorithmic approaches: linear regularization, ensemble learning, and gradient boosting, respectively, providing a robust baseline for comparison. To ensure a robust evaluation that accounts for both the relative fit and the clinical relevance of the predictions, we employed a multi-dimensional assessment strategy. Each method’s predictive performance was assessed using the Coefficient of Determination (*R*^2^), the Root Mean Square Error (*RMSE*), and the Ratio of Performance to Deviation 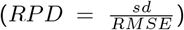 . These metrics offer complementary perspectives on the algorithm fit: *R*^2^ quantifies the proportion of variance (signal) captured, RMSE provides a measure of absolute error in the original scale of the dependent variable, which is essential for determining the clinical relevance of the prediction, and RPD normalizes the RMSE by the standard deviation of the target variable, facilitating comparisons across different datasets and cohorts. Each of the selected methods allows for the extraction of important features during the prediction process, defined as the algorithm’s “biomarker”. We evaluated the relevance of these biomarkers based on the availability of the ground truth. For synthetic data with known ground truth, we calculated the Jaccard score [22] (*J*) to measure the overlap between the biomarker set (*B*) and the true feature set (*T*):

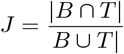

To further characterize the selection errors, we computed the Jaccard-based False Positive (*FPR*_*J*_) and False Negative (*FNR*_*J*_) ratios:

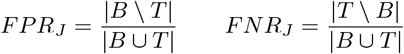

For real-world clinical datasets, where the ground truth is unknown, biomarker quality was assessed through a functional characterization of the features in each biomarker, as detailed in Section 2.7. Extracting a discrete biomarker set is straightforward for BI-SCUDO Regression and LASSO: for the former, it is the union of all sample-specific signatures; for the latter, it comprises all features with non-zero coefficients.

In contrast, Random Forest and XGBoost assign continuous and positive importance scores to all features, making the selection of a cutoff point inherently subjective. To eliminate manual bias, we employed the Kneedle algorithm [23] to perform automated knee-point detection on the ranked feature importance curves computed according to the Mean Decreased Impurity (Gini) approach [4]. This data-driven approach identifies the optimal threshold where the marginal gain in importance drops significantly, ensuring a fair and objective comparison of the biomarkers extracted by each method.

### Synthetic datasets generation

To evaluate the performance and noise resilience of the method, we generated a set of synthetic datasets designed to emulate the characteristics of microarray assays in a controlled environment [24] (we refer to the test on the TCGA datasets for an evaluation of BI-SCUDO on NGS data). The generation process follows four primary steps: metadata simulation, feature profile construction, noise injection, and intensity transformation. We simulated a single continuous dependent variable (the “metadata”) by sampling from a normal distribution:

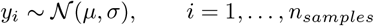

This yields the target vector 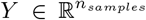, representing the biological or clinical variable to be predicted. Features were generated based on three distinct relationship types with the metadata: correlated, anti-correlated, and non-correlated [25]. The correlated and anti-correlated features constituted the true feature set *T* for Jaccard score computation. For each feature *j*, we first established a maximum expression potential 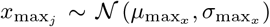 . Each correlated feature was generated as a linear function of the metadata, constrained between 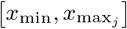:

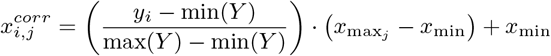

Each anti-correlated feature was generated by producing a correlated feature and then inverting its range:

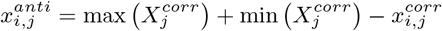

Non-correlated features represent stochastic background expression, and thus were produced by sampling values independently of the metadata:

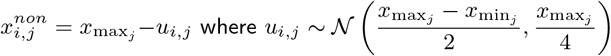

To simulate experimental variability and the noisy nature of biological data, we generated a noise matrix 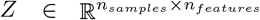, where *z*_*i,j*_ ∈ *Z* is sampled from a feature-specific normal distribution *N* (*µ*_*j*_, *σ*_*j*_), where *µ*_*j*_ and *σ*_*j*_ where both sampled from a standard normal distribution. Noise was added to the signal matrix *X* by using a scaling intensity parameter *a*:

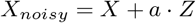

Since microarray signal intensities typically follow an exponential distribution, the final simulated data were transformed to the intensity scale [26]:

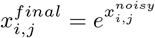

For each in silico experiment, an independent validation dataset was constructed using the same procedure. We resampled the metadata *Y* and noise *Z* while maintaining the feature-specific parameter *x*_*j*_ to ensure the underlying signal remained consistent across the training and testing phases. To assess the predictive performance and noise resilience of the proposed method, we generated 10 datasets independently, each consisting of 1000 features and 150 samples for the training set and 150 for the validation set. For these, we fixed *y* ∼ *N* (120, 40) to obtain enough variability in the metadata to challenge the methods and 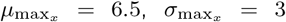 and *x*_min_ = 3.5 to heuristically mimic a typical real world microarray dataset. Finally, we varied the noise factor *a* from 0 to 2, establishing the upper bound at the point where the relative noise level 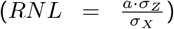, became greater than 1. This happened at *a* = 2. The impact of noise on these datasets is visualized in Figure 1. Additionally, to benchmark the computational efficiency and scalability of BI-SCUDO Regression, we generated high-dimensional datasets containing more than 70000 features. These datasets were simulated with varying sample sizes (*n* ∈ *{*100, 150, 500, 1000*}*), noise level *a* = 0.5, and keeping the other parameters at the same values as before to measure the impact of cohort size on algorithm runtime. Computational benchmarks were performed on a high-performance computing (HPC) cluster using 40 allocated cores. The hardware environment consisted of nodes equipped with Intel Xeon E5-2650V3 (10-core, 2.30 GHz), E5-2680 v4 (14-core, 2.40 GHz), or Gold 6252N (24-core, 2.30 GHz) processors.

**Figure 1.**
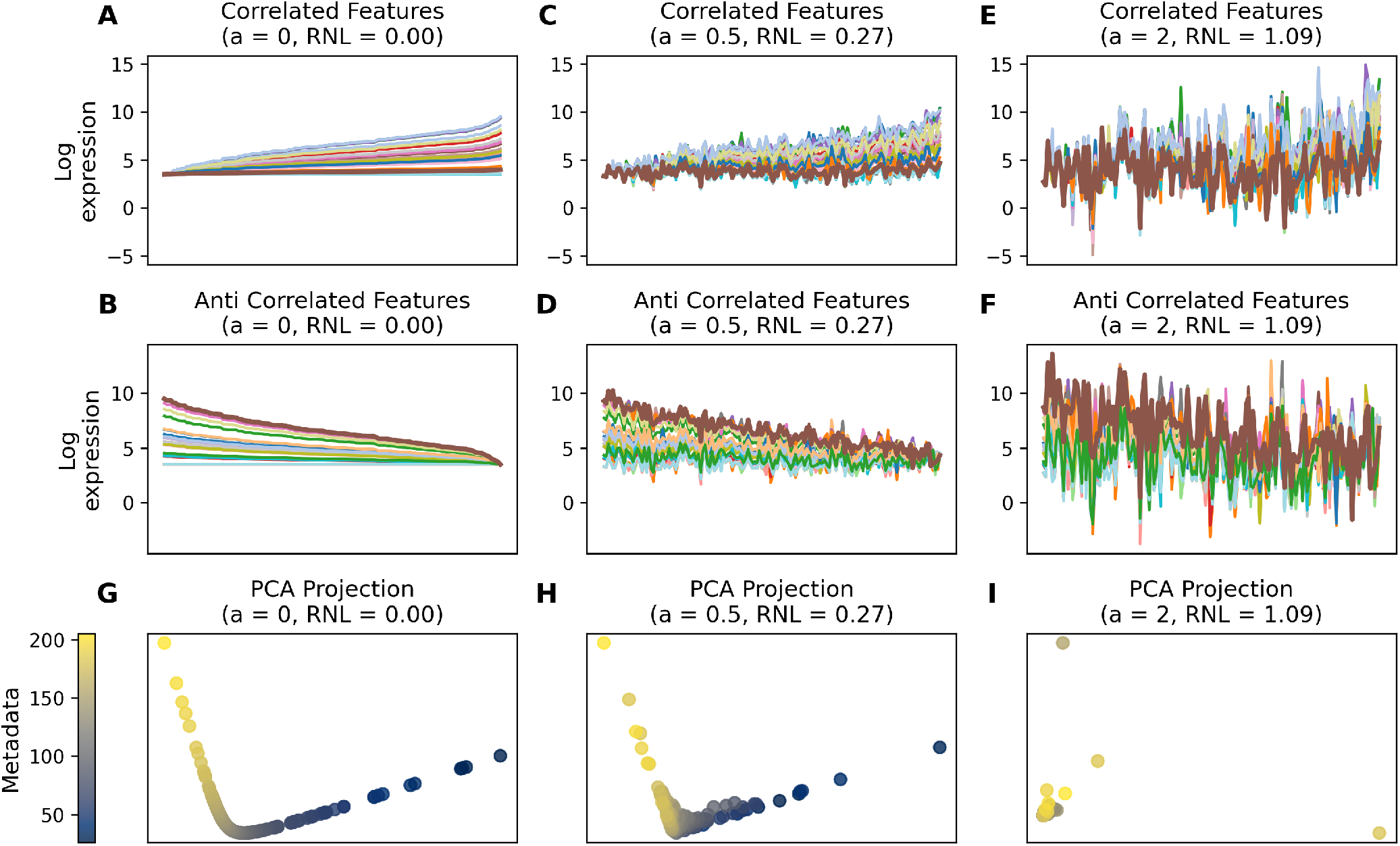
Impact of Additive Noise on Dataset Structure and PCA Manifold. **(A-F)**: Log-scale expression profiles of simulated features. Top row: Features positively correlated with metadata. Middle row: Features anti-correlated with metadata. Noise scaling amplitudes increase from left to right: **(A, B)** *a* = 0 (*RNL* = 0.0), **(C, D)** *a* = 0.5 (*RNL* = 0.25), and **(E, F)** *a* = 2 (*RNL* = 1.1), revealing the progressive obscuration of the underlying biological signal. **(G-I)** Principal Component Analysis (PCA) projections on the full datasets at corresponding noise levels. Each point is a sample, colored according to its continuous simulated metadata value (see color bar). The progression from a distinct, structured manifold ordering **(G)** to a randomized cloud **(I)** visualizes the signal degradation that feature selection and regression methods must overcome.

### Accessing data from The Cancer Genome Atlas

We retrieved transcriptomic and clinical data from The Cancer Genome Atlas (TCGA) [20] using an automated pipeline based on the TCGAbiolinks R library [27]. We downloaded all public projects containing transcriptomic profiles and clinical metadata. Raw sequencing counts were converted to Transcripts Per Million (TPM) [28] to account for transcript length and sequencing depth and log-transformed. Consistent with the design of BI-SCUDO Regression, which leverages the rank-based properties of gene expression within individual samples, no further cross-sample normalization or batch correction was applied. This approach preserves the relative internal structure of the signatures, central to the algorithm’s “normalization-free” logic. To ensure the target variables, overall survival (*OS*) and disease-specific survival (*DSS*) were continuous and fully observed, we restricted the analysis to deceased patients. Each cancer project was then stratified into four distinct experimental datasets based on the AJCC pathological stage following information from [29]: (i) No-stage: all samples regardless of pathological stage, (ii) I-IV: samples spanning the full clinical progression, (iii) II-IV: exclusion of early stage samples, and (iv) III-IV: focus on late stage samples. Only datasets containing more than 30 samples were retained for analysis. This filtering process yielded 39 datasets for OS and 32 for DSS, comprising 3455 and 2369 distinct samples, respectively, which served as the basis for evaluating BI-SCUDO Regression, LASSO, Random Forest Regression, and XGBoost via 10-fold cross-validation. Detailed metrics regarding sample sizes and survival variability across projects are detailed in Figure 2 and Figure 3. To evaluate the sensitivity of BI-SCUDO Regression to batch effects, PERMANOVA *R*^2^ [30] was computed across sample plate IDs. Analysis was performed on three distinct distance matrices: (i) uncorrected Euclidean distance, (ii) Euclidean distance following ComBat batch correction [31], and (iii) the distance matrix derived by BI-SCUDO Regression.

**Figure 2.**
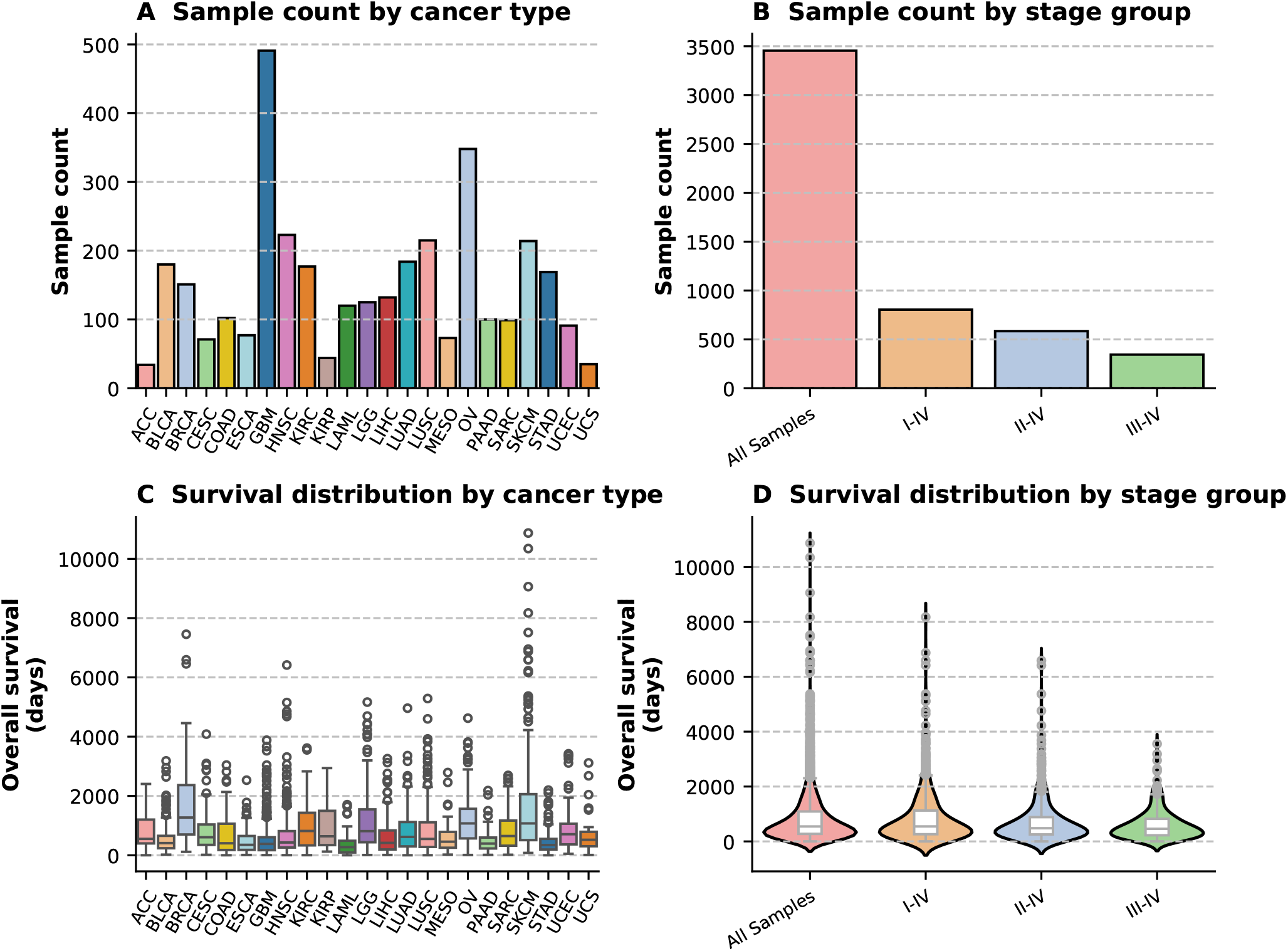
Composition and survival characteristics of the processed TCGA cohorts (overall survival). **(A)** Total sample count per cancer project after filtering for deceased patients. **(B)** Average sample size (*±σ*) across the four disease-stage stratification tiers, showing the expected decrease in cohort size as staging criteria become more restrictive. **(C)** Distribution of overall survival (*OS*) times across cancer projects. Boxplots indicate the median and interquartile range, with individual points representing out-of-range patients. **(D)** Overall survival distribution stratified by the four experimental dataset tiers. Note the high variability in survival times across all projects and stages, and how it gets slightly lower as the staging criteria become more restrictive.

**Figure 3.**
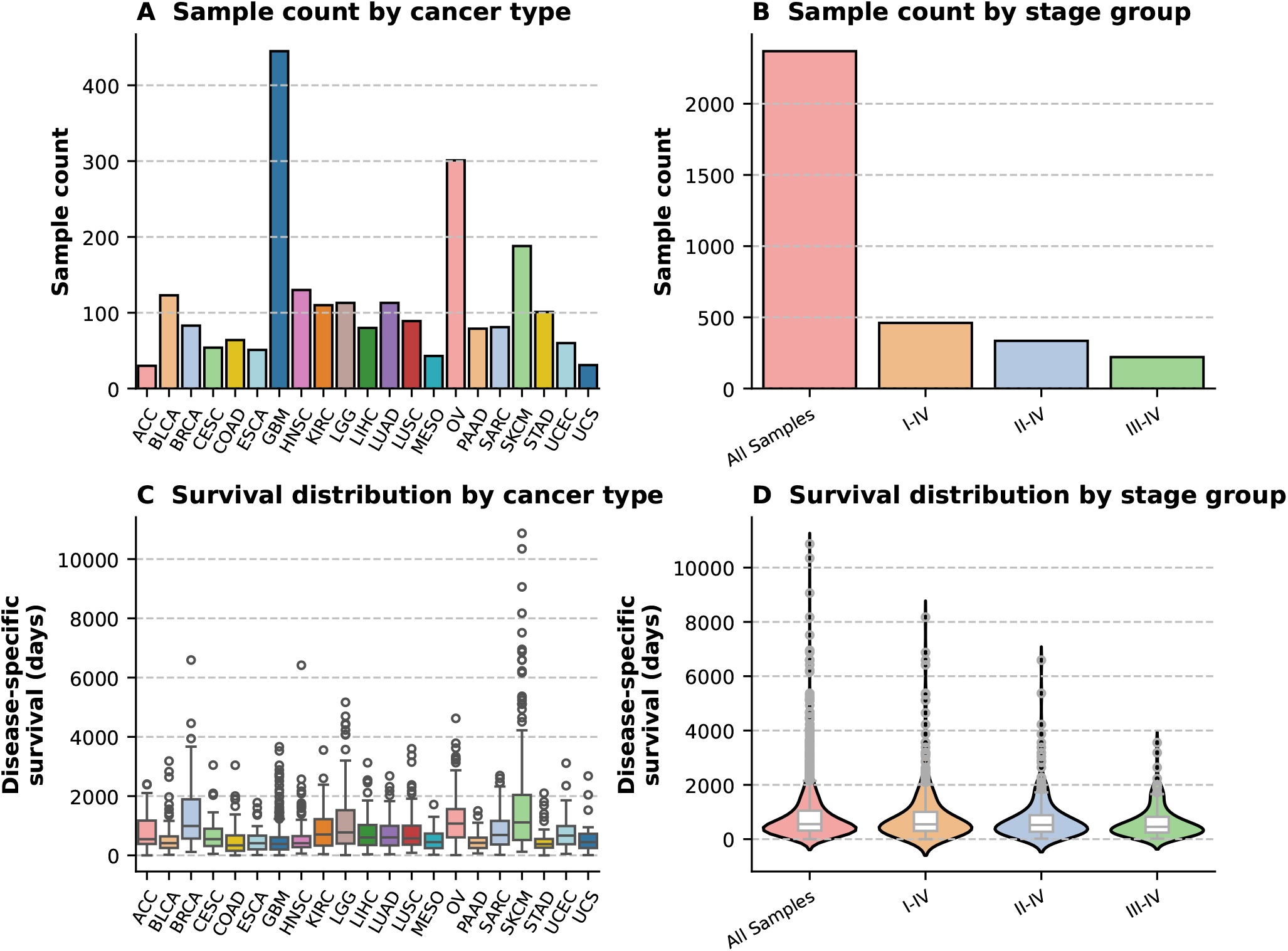
Composition and survival characteristics of the processed TCGA cohorts (disease specific survival). **(A)** Total sample count per cancer project after filtering for deceased patients. **(B)** Average sample size (*±σ*) across the four disease-stage stratification tiers, showing the expected decrease in cohort size as staging criteria become more restrictive. **(C)** Distribution of overall survival (*DSS*) times across cancer projects. Boxplots indicate the median and interquartile range, with individual points representing out-of-range patients. **(D)** Overall survival distribution stratified by the four experimental dataset tiers. Note the high variability in survival times across all projects and stages, and how it gets slightly lower as the staging criteria become more restrictive.

### Functional Enrichment and Biomarker Validation

To evaluate the biological relevance of the biomarkers extracted from the TCGA datasets, we developed a functional annotation pipeline to categorize features as either cancer-related or not. Since a definitive “ground truth” is unavailable for real-world clinical data, we utilized a keyword-based interrogation of four major pathway databases: Gene Ontology Biological Processes (GO:BP) [32] via the GO.db R library, KEGG Pathways [33] via KEGGREST, Reactome [34] via ReactomePA and the MsigDB [35] via msigdbr.

We retrieved all patway description for every Ensembl ID present in the TCGA training sets. A gene was tagged as cancer-related if any associated pathway containd at least one oncology-specific keyworkd, in particular: *cancer, carcinoma, tumor, tumour, oncogen, malign, neoplasm, metasta, leukemia, leukaemia, lymphoma, sarcoma, melanoma, glioma, blastoma, adenocarcinoma, carcinogenesis, tumorigenesis* or *oncogenic*. Of the 20439 unique Ensemble IDs identified across the TCGA cohorts, 1152 (5.6%) were categorized as cancer-related through this procedure. We assessed the quality of the biomarkers extracted by BI-SCUDO Regression, LASSO, Random Forest Regression and XGBoost based on the enrichment of these cancer-related genes. For each extracted biomarker, we counted the number of cancer-related genes appearing in it and performed a one-sided Fisher’s exact test [36] to determine if the proportion of cancer-related genes was significantly higher than would be expected by chance. Biomarkers yielding a p-value *<* 0.05 were considered to contain a statistically significant enrichment of biologically relevant information.

## Results

### BI-SCUDO Regression: an explainable, rank-based regression method for omics data

The need for an explainable regression method capable of handling high-dimensional omics data led to the development of the BI-SCUDO Regression framework. The method was implemented in Python 3.14. The core architecture of the algorithm, outlined in Figure 4, is driven by a six-step procedure designed to extract robust, patient-specific prognostic signatures.

**Figure 4.**
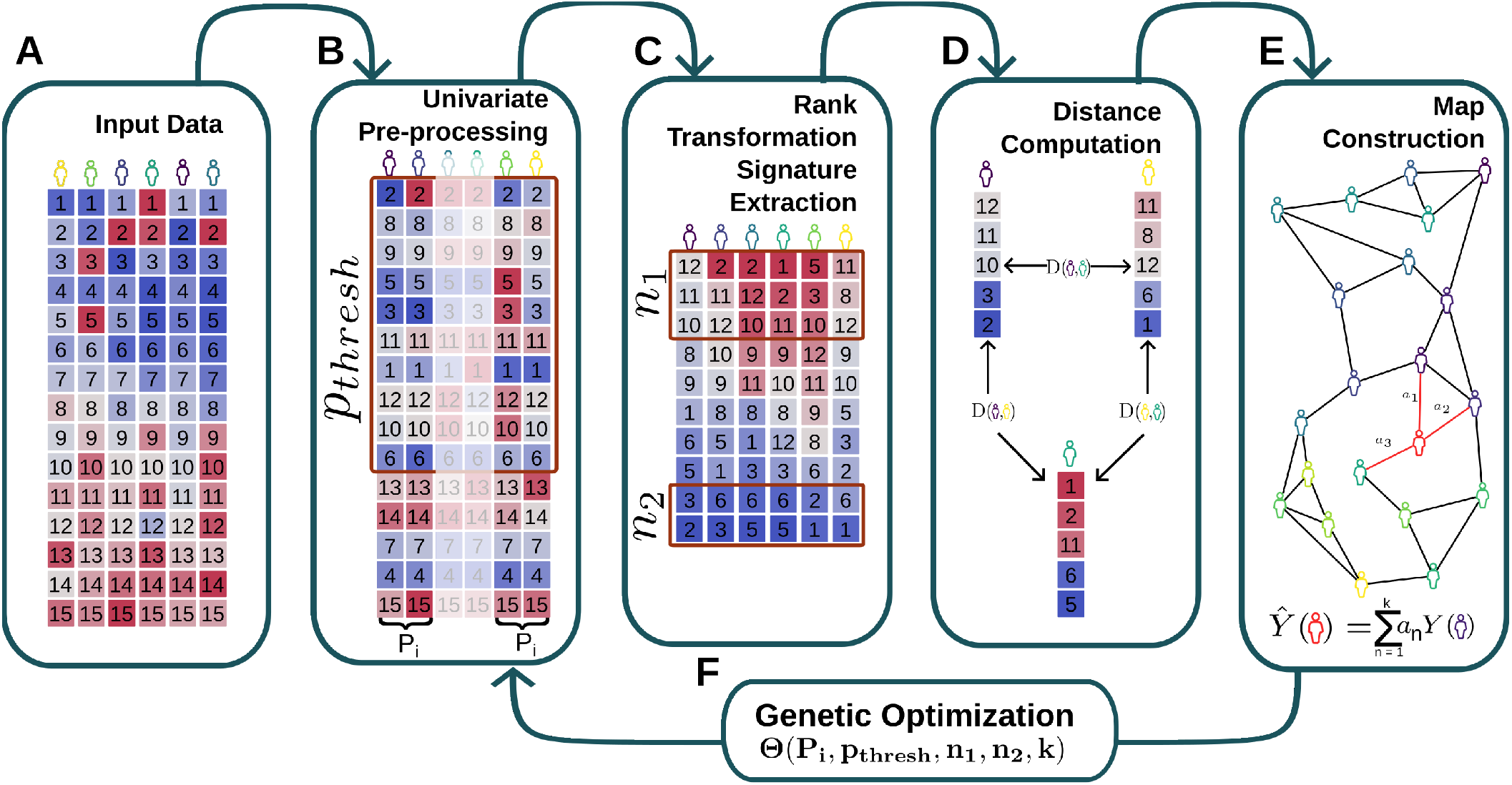
Schematic workflow of the BI-SCUDO regression algorithm. **(A) Input Data:** The initial high-dimensional molecular dataset across multiple samples/patients. **(B) Univariate Pre-processing:** Initial feature selection step governed by a significance threshold (*p*_*thresh*_) obtained from a Wilcoxon rank-sum test on the top and bottom *P*_*i*_-percentiles. **(C) Rank Transformation and Signature Extraction:** Transformation of data into ranks to extract patient-specific signatures composed of the top *n*_1_ and bottom *n*_2_ features. **(D) Distance Computation:** Calculation of pairwise distance metrics between the extracted sample signatures. **(E) Map Construction:** Building a sample similarity network with edges between a sample and its neighbors based on pairwise distances to execute localized regression-based predictions. **(F) Genetic Optimization:** An evolutionary feedback loop that iteratively optimizes the five core hyperparameters Θ(*P*_*i*_, *p*_*thresh*_, *n*_1_, *n*_2_, *k*) to maximize regression performance.

Given a training cohort, BI-SCUDO Regression first divides the samples into “top” and “bottom” proxy classes based on the extreme *P*_*i*_ percentiles of the continuous response variable distribution. A univariate Wilcoxon rank-sum test (Figure 4 Panel B) is applied to each feature to assess its differential expression between these extremes. A dynamic threshold (*p*_*thresh*_) is defined, and features with *p*-values exceeding this limit are discarded. This critical filtering step removes variables that lack response variable-dependent variation, such as housekeeping genes or stochastic noise, performing a first dimensionality reduction step. The expression values of the retained features are subsequently transformed into within-sample rankings. Although this transformation slightly reduces quantitative resolution, it drastically increases the model’s resilience to technical noise and inter-batch variability without the need for cross-sample normalization [37]. For each sample, the *n*_1_ highest-ranked and *n*_2_ lowest-ranked features are extracted, forming a patient-specific signature of length *n*1 + *n*2 (Figure 4 Panel C).

To map the topology of the patient cohort, pairwise distances between samples are computed as a function of the overlap and structural similarity between their signatures, utilizing a weighted enrichment score metric [21] (Figure 4 Panel D). A directed graph is then constructed where nodes represent samples, and edges connect each node to its *k*-nearest neighbors. The neighborhood size is constrained during the optimization procedure (3 *≤ k ≤* 15) because the weighted enrichment score is optimized to detect strong structural overlaps. Restricting connections to the most similar samples mitigates the noise introduced by computing distances between divergent samples. The procedure is detailed in Section 2.1 (Figure 4 Panel E).

To predict the continuous target variable for a new sample, its rank-based signature is extracted and queried against the established training network. The new sample is embedded into the graph, its nearest neighbors are identified, and the predicted value is computed as the distance-weighted average of those neighbors’ response variable values. Notably, the pairwise distance matrix generated during training can also be projected via Principal Coordinate Analysis (PCoA) [38], facilitating direct visualization of the dataset’s manifold and providing a qualitative assessment of the regression structure (Supplementary Figure S1).

Because there is no analytical ground truth for selecting the optimal hyperparameters Θ(*P*_*i*_, *p*_*thresh*_, *n*_1_, *n*_2_, *k*), these values are determined automatically using a custom Genetic Algorithm (GA) implemented in Python via the DEAP [39] framework (Figure 4 Panel F). Model fitness is evaluated within a cross-validation framework (leave-one-out by default) to minimize predictive error. To ensure efficient convergence toward a valid parameter space, the optimization incorporates: (i) rejection sampling and constraint-repair mechanisms to enforce minimum signature length and feature availability bounds, (ii) exponential mutation step-size (*σ*) decay to transition from exploration to exploitation, and (iii) elitism combined with duplicate culling to maintain population diversity and prevent premature convergence. The objective function minimizes the Root Mean Square Error (RMSE) while imposing a penalty proportional to the signature length, thereby favoring parsimonious, highly interpretable models. Complete GA configurations are reported in Supplementary Table S1.

Upon completion, the framework returns the optimized predictive model, cross-validated performance metrics, the discrete patient-specific signatures, the patient distance matrix, and a final consensus biomarker, defined as the union of all features included in the training signatures.

To rigorously validate the statistical significance of the optimized model and ensure the framework did not overfit the high-dimensional data, we implemented a permutation testing procedure based on within-sample feature randomization. In this approach, the expression values of the features are permuted independently within each individual subject. This randomization intentionally destroys the intra-sample biological pathways and the specific relative rankings of the genes, which form the core of the BI-SCUDO feature signature, while perfectly preserving the original clinical response variable distribution and the global expression intensity profile of each sample. The complete predictive pipeline, including the distance matrix computation and neighbor embedding, is executed on this randomized data for *N* iterations (*N* = 10000). For each permutation, the predictive performance is quantified using the Ratio of Performance to Deviation (*RPD*). An empirical *p*-value is subsequently calculated as the fraction of permutation runs that achieve an *RPD* equal to or greater than the *RPD* of the true optimized model. Although the absolute nature of *RMSE* makes it an ideal objective function for the genetic algorithm, *RPD* was chosen for final validation because it is dimensionless and scale-independent. Crucially, because it scales inversely and monotonically with RMSE, RPD provides a mathematically consistent yet highly intuitive standard for external reporting. This validation step guarantees that the predictive power of the BI-SCUDO Regression biomarker stems from genuine, specific transcriptomic patterns rather than algorithmic artifacts, data leakage, or stochastic noise [40], [41].

### BI-SCUDO regression outperforms LASSO, Random Forest Regression and XGBoost in noisy in-silico datasets

BI-SCUDO Regression was first benchmarked against three state-of-the-art regression methods: LASSO, Random Forest Regression and XGBoost, using the fully in-silico setup described in Section 2.5.

The global performance metrics *R*^2^, *RMSE* and *RPD* obtained from 10 different datasets reveal distinct profiles for each algorithm under increasing noise, as seen in Figure 5.

**Figure 5.**
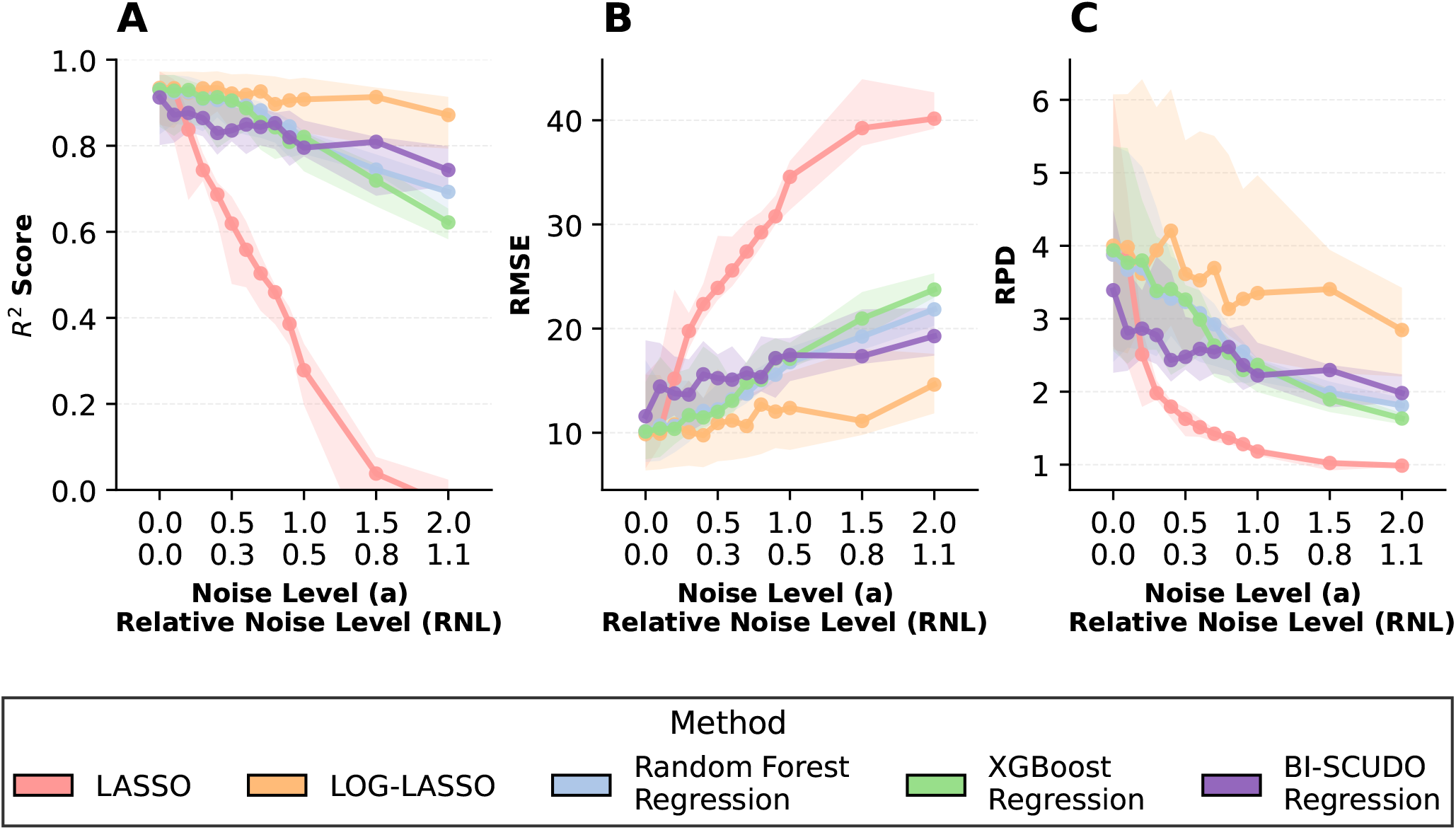
Predictive performance of regression models under varying degrees of stochastic noise. The panel displays the **(A)** *R*^2^ scores, **(B)** Root Mean Square Error (*RMSE*), and **(C)** Ratio of Performance to Deviation (*RPD*) for LASSO, LASSO applied to log-transformed data, Random Forest Regression, XGBoost and BI-SCUDO Regression. Evaluations were performed across 10 independent synthetic datasets (150 samples, 1000 features), subjected to white noise with an amplitude ranging from 0 to 2.0, obtaining a relative noise level increasing from 0 to 1.1. Dots represent the median value of the 10 runs with the corresponding interquartile range band. While all models perform similarly with zero noise, BI-SCUDO Regression demonstrates superior resilience and maintains the highest predictive capacity at extreme noise levels. LOG-LASSO represents the upper bound of predictive performance, as the log-transformation successfully restores the underlying linear relationship between metadata and feature values.

LOG-LASSO serves as an indication of the ideal benchmark upper bound across all noise levels, as the log-transformation successfully restores the true linear relationship within the synthetic data (*R*^2^ = 0.934 and *RPD* = 4.00 at *RNL* = 0.0, declining only slightly to *R*^2^ = 0.872 and *RPD* = 2.85 at extreme noise *RNL* = 1.1). At the baseline, all models perform close to this theoretical limit (median *R*^2^ *>* 0.91, *RPD >* 3.39). The first divergence point occurs at *RNL* = 0.11, where the LASSO regressor begins a sharp collapse: its median *R*^2^ falls to 0.84 and median RPD to 2.51, while the other methods maintain a median *R*^2^ ≥ 0.88 (LOG-LASSO *R*^2^ = 0.92) and *RPD >* 2.86. Following this first threshold, LASSO performances degrade rapidly and by *RNL* = 0.81 it becomes non-predictive (median *R*^2^ = 0.038 and *RPD* = 1.02) and collapsing completely at maximum noise (*RNL* = 1.1, median *R*^2^ = *−*0.035 and *RPD* = 0.99). At low-to-moderate noise levels (*RNL ≤* 0.16), Random Forest Regression and XGBoost Regression perform closest to the LOG-LASSO upper bound (*R*^2^ ∈ [0.91, 0.93] and *RPD* ∈ [3.36, 3.94]) compared to BI-SCUDO Regression (*R*^2^ ∈ [0.86, 0.87] and *RPD* ∈ [2.78, 3.40]). This performance gap stems from the loss of quantitative resolution inherent to BI-SCUDO’s rank-transformation step; the relative rank order acts as a quantization bottleneck that removes signal when the data is clean. However, as noise increases (*RNL ≥* 0.43), this design choice turns into BI-SCUDO’s primary asset. Heavy noise severely corrupts continuous feature magnitudes, causing XGBoost and Random Forest Regression to overfit to random fluctuations and experience rapid performance degradation (*R*^2^ = 0.62 and 0.69 and *RPD* = 1.63 and 1.81 at *RNL* = 1.1 respectively). In contrast, ordinal rank ordering functions as a robust topological filter that remains invariant to additive amplitude perturbations. Consequently, BI-SCUDO trades away minor baseline resolution in pristine data to achieve unmatched stability under heavy noise conditions (*R*^2^ = 0.74 and *RDP* = 1.98 at *RNL* = 1.1). The ability to recover true features correlated with the dependent variable was assessed through the Jaccard score (*J*), false positive ratio (*FPR*_*J*_), and false negative ratio (*FNR*_*J*_), as shown in Figure 6.

**Figure 6.**
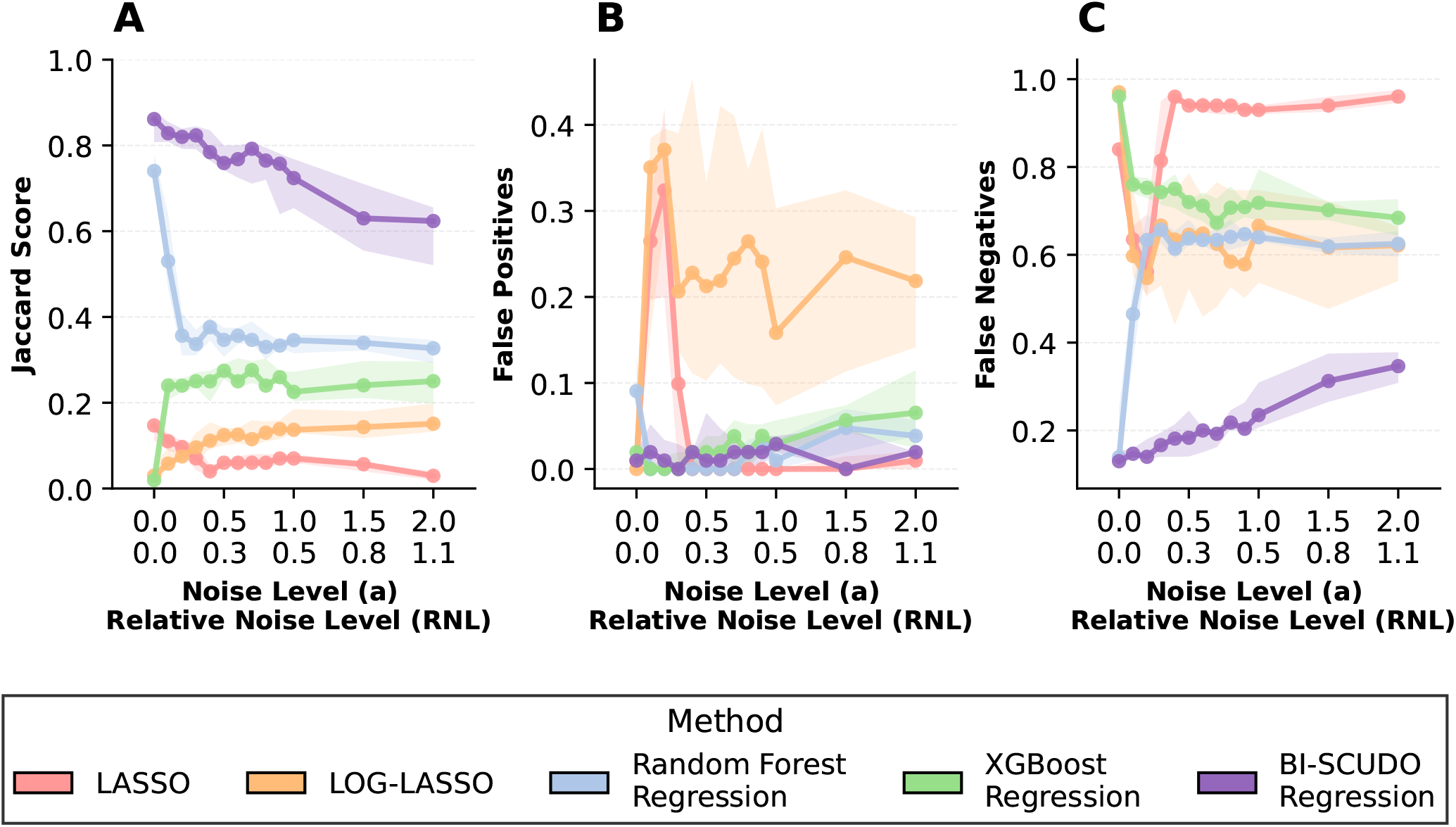
Feature selection accuracy across increasing noise levels. The panels illustrate the **(A)** Jaccard Index, **(B)** False Positive ratio, and **(C)** False Negative ratio for the feature sets extracted by Lasso, Log-LASSO, Random Forest, XGBoost, and BI-SCUDO Regression compared against the features correlated with the dependent variable. Dots represent the median value of the 10 runs with the corresponding interquartile range band. BI-SCUDO maintains the highest structural similarity to the true biomarker signature across the entire noise spectrum, driven by a low False Positive ratio and a lower False Negative ratio compared to the other algorithms.

A stark divergence in feature selection capability is visible even at baseline (*RNL* = 0.0), where BI-SCUDO Regression achieves a median Jaccard score of 0.861, whereas standard LASSO (*J* = 0.147), LOG-LASSO (*J* = 0.030), and XGBoost (*J* = 0.020) fail to isolate the true signature. Standard LASSO suffers from severe feature omission due to collinearity, exhibiting an *FNR*_*J*_ of 0.635 already at *RNL* = 0.05 and exceeding 0.90 for *RNL ≥* 0.27 (despite keeping *FPR*_*J*_ *≤* 0.02). XGBoost similarly suffers from high omission rates (*FNR*_*J*_ *≥* 0.68) across all noise levels. Random Forest Regression exhibits a rapid decline in feature recovery early on, with its Jaccard score falling from 0.740 at *RNL* = 0.0 to 0.357 at *RNL* = 0.11 as its false negative rate climbs to 0.634. BI-SCUDO Regression significantly outperforms all benchmark methods in biomarker identification, maintaining a high Jaccard score (*J ≥* 0.724) and low false negative rate (*FNR*_*J*_ *≤* 0.235) up to *RNL* = 0.54. Even under maximum noise (*RNL* = 1.1), BI-SCUDO Regression retains a remarkably strong Jaccard index of 0.624 paired with near-zero false positives (*FPR*_*J*_ = 0.020). This highlights the effectiveness of the Genetic Algorithm’s parsimony penalty in enforcing exceptional specificity even in high-noise environments.

The permutation test on the BI-SCUDO Regression biomarkers resulted in a *p*-value of less than 10^*−*4^ in all synthetic runs. This ensures that every biomarker identified by the BI-SCUDO Regression algorithm isolates high-confidence signals even when they are heavily masked by stochastic noise.

Finally, the computational overhead necessary to train BI-SCUDO Regression on a dataset was evaluated using 10 independently generated datasets with more than 70, 000 features and varying sample sizes (100, 150, 500, 1000) generated according to Section 2.5 (Figure 7). At a low sample count, BI-SCUDO demonstrates a competitive median execution time, closely matching LASSO, XGBoost, and

**Figure 7.**
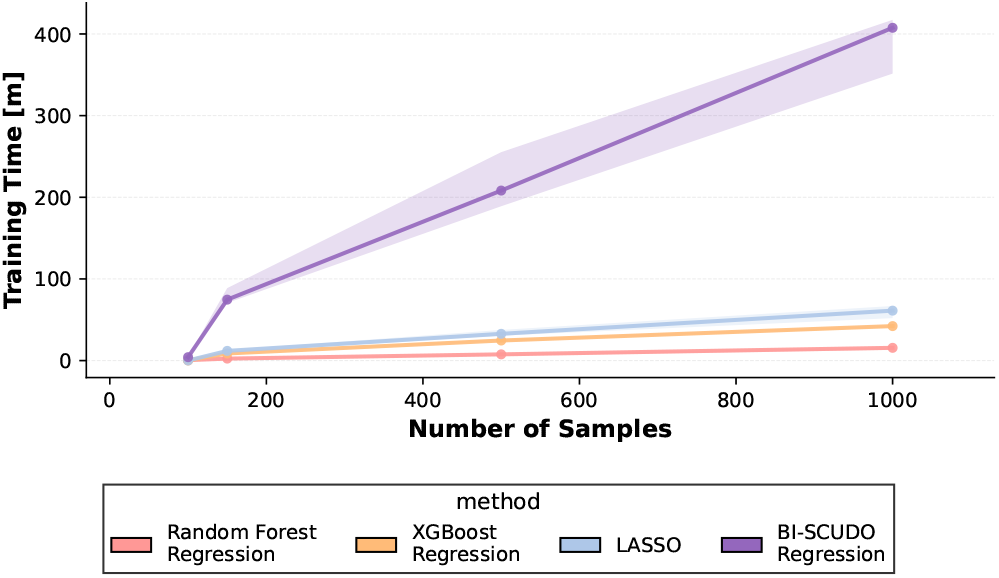
Training time as a function of sample size. Time (in minutes) required to train BI-SCUDO Regression, LASSO, Random Forest Regression, and XGBoost on datasets with varying numbers of samples.

Random Forest Regression. However, as the sample size increases, the computational cost of BI-SCUDO Regression scales steeply due to the pairwise distance matrix construction and the iterative search of the Genetic Algorithm. At 1000 samples, it requires around 6.7 hours of computation time, while LASSO, XGBoost regression, and Random Forest Regression require around 1 hour, 42, and 15 minutes, respectively.

### BI-SCUDO Regression outperforms LASSO, Random Forest Regression and XGBoost in predicting overall survival across the TCGA database

To assess the applicability of the method in real-world precision medicine scenarios, we performed regression analysis on overall and disease-specific survival from cancer patients in the TCGA database. Project filtering and stratification are detailed in Section 2.6.

The application of BI-SCUDO Regression to high-dimensional transcriptomics survival data reveals a robust capacity to extract signal where traditional machine learning architectures largely fail. Performances can be visualized in Figure 8. While LASSO, Random Forest Regression, and XGBoost consistently yielded negative median *R*^2^ when trained across all clinical stratifications for both Overall Survival (*OS*) and Disease Specific Survival (*DSS*), BI-SCUDO Regression maintained stable and positive predictive performances, with *R*^2^ values varying between 0.37 and 0.56 for *OS* and 0.38 and 0.62 for *DSS*. This distinction is most apparent in the absolute error metrics: for instance, in the *No Stage* cohort, BI-SCUDO Regression achieved a median *RMSE* of 361 for *DSS*, representing an approximate 32% reduction in prediction error compared to the *RMSE* = 531 recorded for LASSO, 545 for XGBoost, and 580 for Random Forest Regression. This suggests that BI-SCUDO Regression is particularly suited to the low signal-to-noise ratios inherent in cancerous survival datasets.

**Figure 8.**
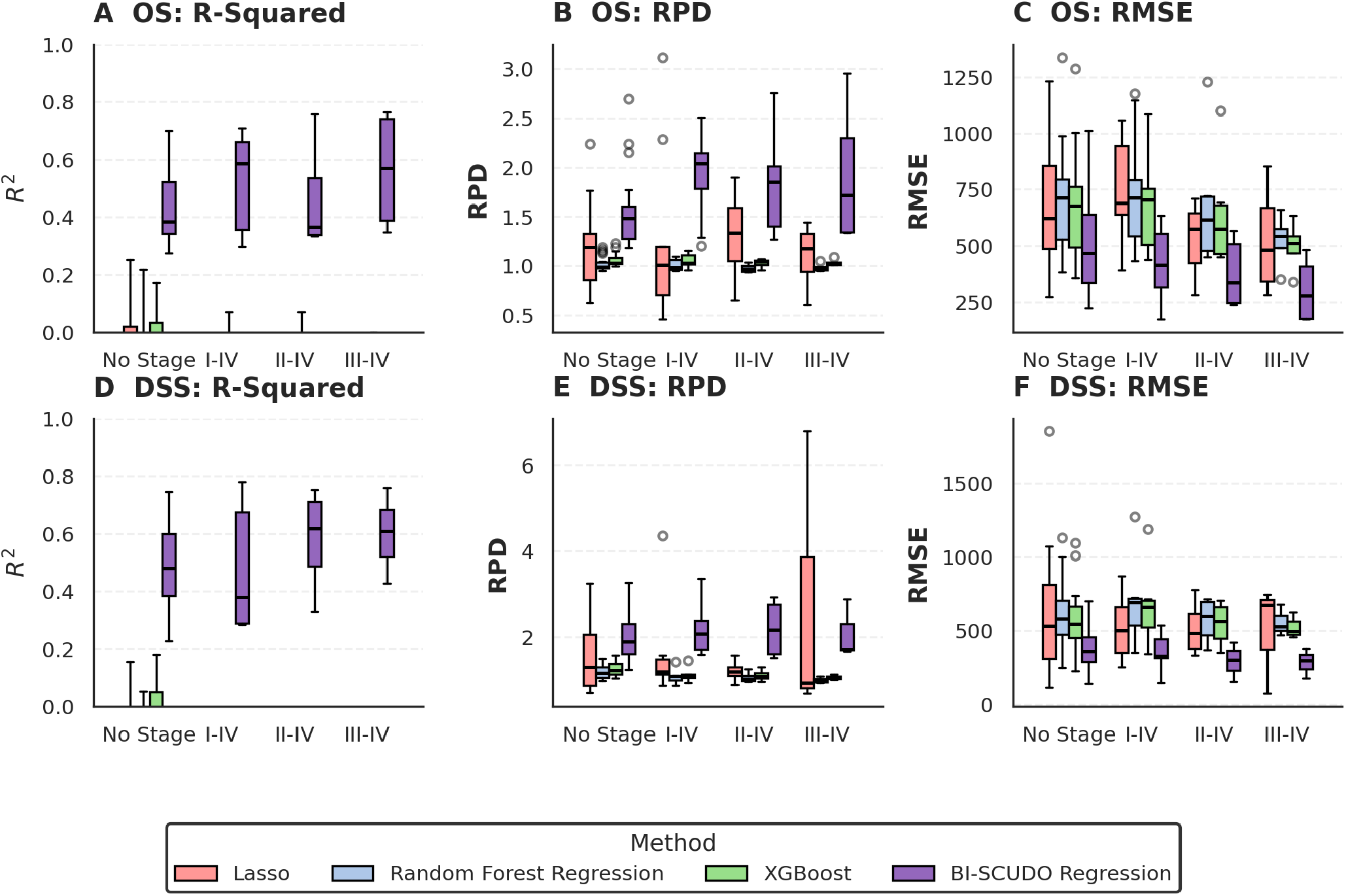
Predictive performance comparison of BI-SCUDO Regression, LASSO, Random Forest Regression, and XGBoost across clinical stratifications. Boxplots detailing the distribution of predictive accuracy metrics across TCGA cohorts, categorized by clinical stage availability (No stage, I-IV, II-IV, and III-IV). Panels display **(A-D)** *R*^2^, **(B-E)** *RPD*, and **(C-F)** *RMSE* for Overall Survival (*OS*) (panels **A, B** and **C**), and for Disease Specific Survival (*DSS*) (panels **D, E** and **F**).

The superiority of BI-SCUDO Regression becomes more pronounced as disease progression advances and biological signals for mortality consolidate. Whereas BI-SCUDO Regression has a mean 34% *RMSE* decrease in *DSS* prediction (31% for *OS*) in the *No Stage* stratification with respect to the state of the art, it reaches a 47% decrease for *DSS* (45% for *OS*) in the late-stage *III-IV* group. This suggests that BI-SCUDO regression is more efficient at capturing critical signal than its competitors, which likely struggle with the reduced sample sizes typical of terminal-stage cohorts.

Beyond numerical accuracy, the biological relevance of the identified biomarkers provides a second layer of validation (Figure 9. In the unstratified analysis (*No Stage*), BI-SCUDO Regression achieved statistically significant enrichment (*p <* 0.05) for known cancer genes in 57.9% (11*/*19) of evaluated *DSS* projects, outperforming XGBoost (47.4%), Random Forest Regression (31.6%), and LASSO (0.0%). This advantage extends across stage-stratified cohorts, where BI-SCUDO reached enrichment rates of 66.7% in Stage *I–IV*, 75.0% in Stage *II–IV*, and 33% in Stages *III-IV*. Internal permutation testing demonstrated robust statistical rigor across the majority of evaluated cohorts, yielding median *p*-values consistently below 0.05 and confirming that BI-SCUDO Regression biomarkers are not products of stochastic noise. Non-significant permutation test results (*p >* 0.05) were restricted to seven specific projects and stratification settings: *TCGA-GBM* (*No Stage, DSS*), *TCGA-HNSC* (*Stage I–IV, DSS*), *TCGA-COAD* (*No Stage, OS*), *TCGA-KIRC* (*Stage I–IV, OS*), *TCGA-KIRP* (*Stage I–IV, OS*), *TCGA-LUAD* (*No Stage, OS*), and *TCGA-SARC* (*No Stage, OS*). Ultimately, BI-SCUDO’s ability to pair high-confidence biological enrichment with superior error mitigation positions it as a significantly reliable tool for transcriptomic survival modeling.

**Figure 9.**
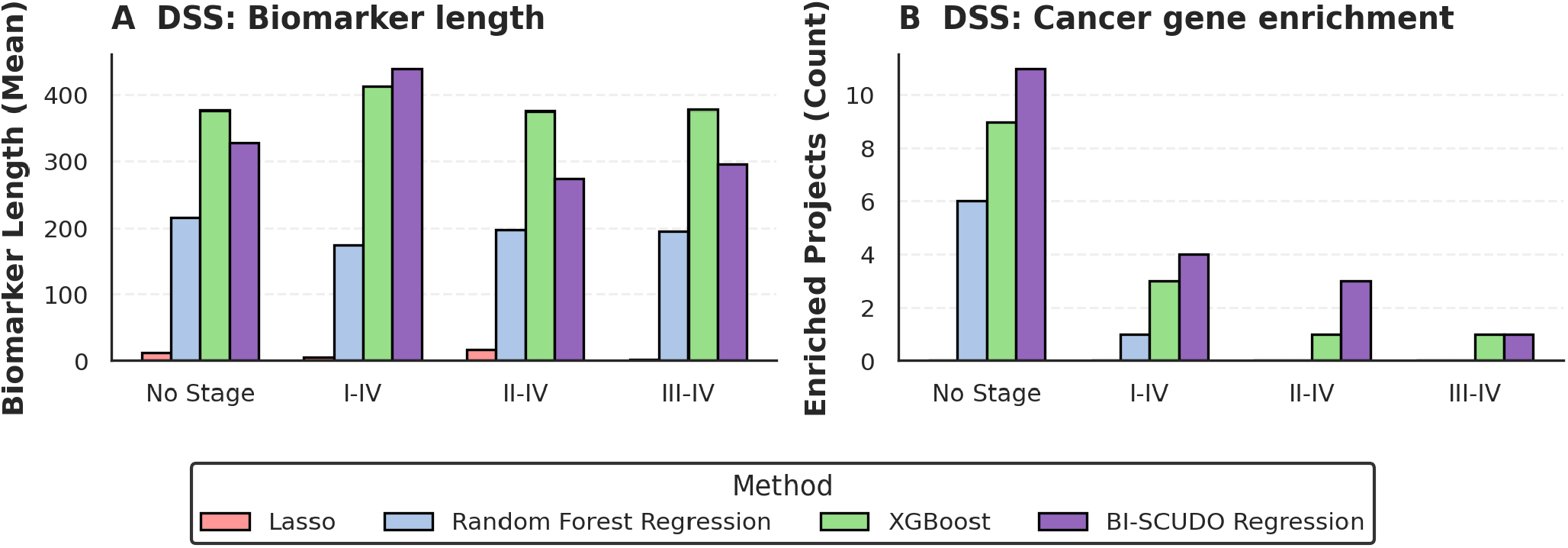
Comparison of BI-SCUDO Regression, LASSO, Random Forest Regression and XGBoost feature selection characteristics and biological relevance of the extracted biomarkers. Analysis of the composition and significance of the gene sets selected by each machine learning method when predicting Disease Specific Survival. **(A)** Bar chart representing the median biomarker length (total number of selected genes) for each method across the four stratifications. **(B)** Bar chart indicating the total number of independent dataset runs that achieved a significant biological enrichment for known cancer drivers after Fisher’s exact test (*p <* 0.05).

To evaluate the impact of technical batch effects, PERMANOVA *R*^2^ was calculated on the distance matrices and the batches detailed in Section 2.6 the methods across stage stratifications and survival measure availability to measure the variance explained by the plate ID batch, where lower values indicate better batch effect mitigation. Across all evaluations, the explicit batch-correction tool ComBat [31] demonstrated the strongest suppression of batch effects, while BI-SCUDO consistently outperformed the uncorrected Baseline by intrinsically reducing batch-driven variance (Figure 10). For Overall Survival (*OS*), ComBat yielded the lowest global median *R*^2^ of 0.0466, followed by BI-SCUDO at 0.1463, whereas the Baseline retained a higher batch variance of 0.2299. A corresponding trend emerged for Disease-Specific Survival (*DSS*), with ComBat achieving a global median of 0.0830, BI-SCUDO reaching 0.1859, and the Baseline measuring 0.2526. As can be assessed through Figure 10, this relative performance hierarchy is maintained consistently across all patient stage stratifications, including *No Stage, I-IV, II-IV*, and *III-IV*. For example, within the *DSS Stage I-IV* cohorts, BI-SCUDO reduced the median *R*^2^ to 0.1999 compared to the Baseline’s 0.2583, though ComBat further reduced it to 0.1172. Similarly, in the *OS* cohorts restricted to *Stages III-IV*, BI-SCUDO (0.1376) mitigated batch effects more effectively than the Baseline (0.2158), trailing ComBat (0.0715). Overall, while BI-SCUDO does not aggressively remove batch variance to the same extent as dedicated harmonization models, it provides a measurable, stable reduction in technical batch effects compared to raw Baseline distances across both survival endpoints and all evaluated stage configurations.

**Figure 10.**
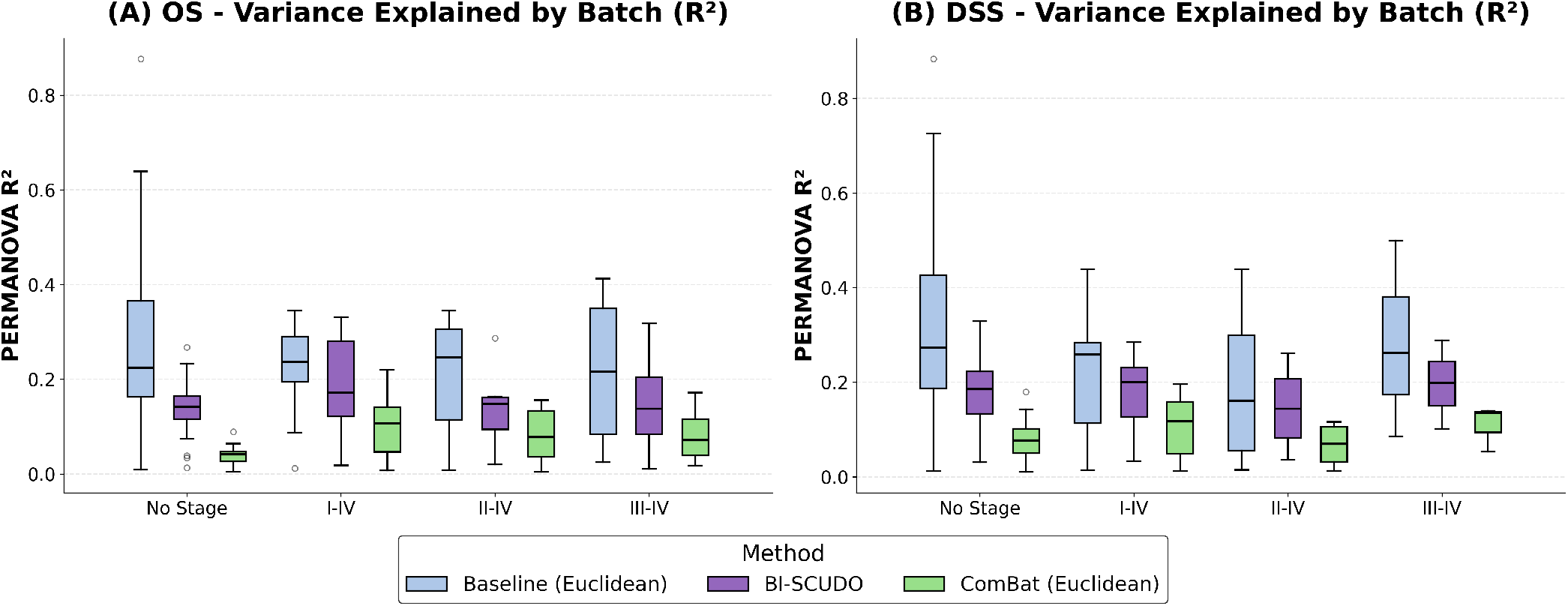
Comparison of batch-effected dependent signal. Permanova *R*^2^ values from plate-id batches between baseline data (light blue), Combat-Normalized Data (green) and BI-SCUDO Regression (purple) distance matrices. **(A)** Overall survival, **(B)** Disease specific survival.

## Discussion

This work introduces BI-SCUDO Regression, a novel rank-based framework designed for the explainable analysis of high-dimensional omics data. By extending a previously validated classification paradigm to continuous outcomes, BI-SCUDO addresses a critical bottleneck in precision medicine: the loss of biological resolution that results from the discretization of clinical variables [42]. While classification-based survival models (e.g., high vs low risk) are useful for broad stratification, they often obscure the granular phenotypic gradients essential for personalized prognostic modeling. The transition to a regression-based architecture allows for a more nuanced interpretation of the transcriptomic landscape, capturing the subtle shifts in gene expression that correlate with disease progression and patient mortality.

At its core, BI-SCUDO Regression leverages within-sample ranks rather than raw expression values. This choice represents a fundamental shift to a nonparametric ordinal space. The primary advantage of rank transformation is its invariance to monotonic transformations. In the context of large-scale meta-analyses and multi-center studies, as in the TCGA test case, batch effects that stem from different sequencing depths, sample preparation protocols, or platform-specific biases often manifest as monotonic shifts in expression intensity [43], [37]. Our empirical evaluation of batch separation highlights the practical benefit of this rank-based transformation. When measuring the variance explained by technical batching (PERMANOVA *R*^2^), BI-SCUDO consistently reduced batch-driven variance compared to uncorrected baseline Euclidean distances across all stage stratifications and survival endpoints. Importantly, BI-SCUDO achieves this mitigation intrinsically: it does not require the explicit batch annotations, covariates necessary for supervised harmonization algorithms to work. While dedicated batch-correction tools that explicitly model and subtract technical variation, like the here tested ComBat [31], can reduce more batch specific variability than BI-SCUDO regression, they rely on pre-defined batch metadata that may be incomplete, unavailable, or confounded with biological variables of interest, which could lead to overfitting and the loss of relevant information. BI-SCUDO’s rank-based architecture serves as an inherent, model-agnostic buffer against platform variance, preserving patient-level relative gene order while naturally attenuating a substantial portion of technical noise without the need to model it explicitly (Supplementary Figure S1). Linear models like LASSO are highly sensitive to these shifts, resulting in coefficients that fail to transfer across cohorts. On the other hand, BI-SCUDO Regression can capture the relative hierarchy of genes that is often more biologically conserved than absolute quantitative levels, providing a molecular fingerprint that remains stable across diverse technical environments [44].

Furthermore, by extracting the “molecular extremes” (*n*_1_, *n*_2_) the framework simultaneously addresses the twin challenges of stochastic noise and high dimensionality. This process filters out the high-variance central part of the expression distribution, where genes are most likely to flip ranking due to stochastic noise, and focuses on the high-confidence signals at the tails. This structural profiling effectively reduces the feature space from tens of thousands of variables to a lean, patient-specific signature. This is particularly relevant in transcriptomics, where the most informative prognostic signal is often concentrated in the genes showing the most dramatic relative dysregulation.

The superior performance of BI-SCUDO on TCGA data highlights a shift in modeling strategy from global linear modeling to local topological manifolds [45]. In fact, traditional regression architectures (LASSO, Random Forest, XGBoost) attempt to fit a single, project-wide rule to the data. This assumes that the relationship between gene expression and survival is uniform across the entire cohort. However, cancer is characterized by extreme inter-patient heterogeneity. Even within a single TCGA project (e.g. *TCGA-BRCA*), the molecular drivers of mortality in a basal-like breast cancer subtype may be entirely different from those in a luminal-A subtype. BI-SCUDO Regression treats the cohort as a topological map rather than a matrix of independent observations. By computing distances through a weighted enrichment score, we measure the structural overlap of signatures, creating a directed graph where edges aim to represent genuine biological kinship. This localized approach acts as a natural filter; a patient’s survival is predicted using only their most similar molecular peers. By segmenting the cohort into small, local neighborhoods, BI-SCUDO effectively learns the complex, multimodal space of overall and disease-specific survival without averaging out distinct biological signals. This explains the framework’s ability to maintain a positive *R*^2^ in projects where global models yielded negative results.

The benchmarking results on synthetic data reveal a significant divergence in noise resilience, which we can attribute to the specific mathematical failure modes of traditional benchmarks when applied to the “small n, large p” problem. Governed by *L*1 regularization, LASSO is designed for sparsity, which is often desirable in clinical settings. However, it is fundamentally greedy when faced with the high collinearity of biological pathways [46]. If fifty genes are correlated within a prognostic pathway, LASSO will select only one representative and zero out the remainder. This can be desirable when looking for clinical biomarkers, but it creates a single point of failure; if stochastic noise corrupts that specific variable in a new sample, the model lacks the redundant biological signal to correct the prediction. Our synthetic results confirm this, showing a total collapse in LASSO’s Jaccard scores as noise amplitude increased beyond *RNL ≥* 0.3. Random Forest Regression, on the other hand, relies on recursive partitioning based on exact expression thresholds. In high-noise scenarios, a noisy split point at the root of a tree propagates errors throughout the entire ensemble [47]. Because survival signals in transcriptomics are often distributed across many genes with subtle effects, Random Forest Regression’s variance-splitting logic frequently misses these distributed signals in favor of a few high-variance features that may simply be artifacts of the training cohort. XGBoost represents a nuanced failure mode: it is the only method with a high false positive ratio in TCGA data, where it finds a number of biomarkers enriched for cancer-related genes similar to the one of BI-SCUDO Regression even when it is unable to make a meaningful prediction [45]. XGBoost appears to overfit to high-variance genes that are widely dysregulated but do not functionally serve as predictors of survival in specific cohorts. This leads to the model memorizing training artifacts rather than capturing the underlying survival manifold, creating biomarkers significantly enriched for cancer-related genes and negative *R*^2^, and in general, poor predictive performances.

A critical distinction in BI-SCUDO’s architecture is its reliance on a Genetic Algorithm for hyperparameter optimization [48]. While ensemble methods such as XGBoost expose a large hyperparameter space (like tree depth, learning rate, number of estimators, and row and column subsampling ratios) whose joint tuning is computationally demanding and prone to overfitting in small cohorts, BI-SCUDO reduces the search space to only five critical variables: *P*_*i*_, *p*_*thresh*_, *n*_1_, *n*_2_, and *k*. By utilizing an evolutionary search to minimize *RMSE* while penalizing signature length, the GA acts as a model architect, enforcing a principle of parsimony. This minimalist approach is essential for stability in the low-sample regimes (*n <* 500) typical of most TCGA projects. The stability of BI-SCUDO’s permutation p-values (*p <* 0.05 in 90% of the different configurations) further confirms that the discovered signatures are products of genuine transcriptomic patterns rather than algorithmic artifacts.

Despite these strengths, several limitations warrant consideration. The transformation to within-sample rankings involves an intentional resolution trade-off. By discarding absolute magnitudes, the model may lose sensitivity to subtle, dosage-dependent effects in genes where expression variance is highly informative. Additionally, the BI-SCUDO Regression framework still presents a scalability challenge. Because graph construction and signature comparison scale quadratically (*O*(*n*^2^)), BI-SCUDO Regression training is currently optimized for clinical cohorts of moderate size (*n <* 2000). While this covers a majority of current precision medicine studies, transitioning to biobank-scale datasets would require the integration of optimized GPU-accelerated distance kernels or approximate nearest-neighbor algorithms to maintain feasibility. Furthermore, the computation time can be reduced by relaxing the convergence criteria for the Genetic Algorithm optimization. Finally, we must emphasize that the signatures identified are associative. While BI-SCUDO Regression consistently isolates gene sets enriched for known cancer drivers, further functional validation is required to distinguish true drivers of mortality from passenger biomarkers.

Looking forward, we aim to extend the BI-SCUDO architecture to multimodal inputs, integrating transcriptomics with DNA methylation, proteomics, and copy number variations [49], [50]. By leveraging synergies across different omic layers, the model could provide a truly holistic view of the patient’s molecular state. Furthermore, we intend to apply this framework to a diverse range of novel, independent datasets to facilitate the discovery of previously uncharacterized molecular drivers of disease.

By pairing high-confidence biological enrichment with superior noise mitigation, BI-SCUDO Regression provides a stable, explainable, and reliable tool for the next generation of precision medicine.

## Conclusion

This study introduces BI-SCUDO Regression, a robust rank-based algorithm developed to handle the unique challenges of high-dimensional omics data. By extending the SCUDO classification paradigm to continuous outcomes, we address a persistent bottleneck in precision medicine: the information loss inherent in the arbitrary discretization of clinical variables. Through the integration of within-sample rank transformations and local topological modeling, BI-SCUDO Regression captures the granular phenotypic gradients that define complex disease states, moving beyond global averages to provide a patient-centric prognostic tool. Our benchmarking results across both synthetic and real-world TCGA datasets demonstrate a significant increase in noise resilience. While traditional linear and tree-based models, such as LASSO, Random Forest, and XGBoost, showed a marked susceptibility to regularization collapse and overfitting in high-noise environments, BI-SCUDO Regression maintained stable predictive accuracy and high feature recovery scores. Its success in the TCGA survival projects, where it was the only method to consistently achieve positive *R*^2^ values, underscores the superiority of local manifold learning over global coefficient-based modeling in the “small n, large p” regime. Furthermore, the ability to assign a statistical significance value to each biomarker introduces another level of control over the model’s performance. In particular, BI-SCUDO Regression was able to extract significant biomarkers in all of the synthetic datasets and in more than 90% of the TCGA datasets.

While the framework is subject to inherent limitations regarding the computational scaling of its *O*(*n*^2^) distance computations and the resolution trade-off associated with ordinal ranking, the model’s unparalleled robustness and interpretability make it an excellent choice to compute predictions and extract meaningful information from complex, low-signal omics datasets. Unlike “black-box” alternatives, BI-SCUDO provides direct visibility into the molecular signatures driving each individual prediction, bridging the gap between mathematical performance and biological insight. In particular, its ability to extract patient-specific signatures makes BI-SCUDO Regression a natural fit for precision medicine applications, where molecular characteristics of single patients or of specific sub-populations need to be easily explored. Ultimately, BI-SCUDO Regression offers a conceptually elegant yet computationally powerful addition to the analytical repertoire of systems biology.

By providing a stable foundation for the discovery of non-linear molecular drivers, this work paves the way for future extensions into multi-omics integration and the identification of novel therapeutic targets. As we transition into an era of increasingly heterogeneous and multi-layered genomic data, BI-SCUDO Regression stands as a reliable framework for the next generation of precision medicine and clinical decision-support systems.

## Data Availability

BI-SCUDO Regression is available at https://github.com/cibio-computational-modeling-lab/Fantoni-et-al-BI-SCUDO-Regression

## Author Contributions Statement

G.F. and L.M. conceived the project, developed its main conceptual ideas, and designed the research plan. G.F. developed the computational framework and performed the computational analyses. M.L., C.A.R., and R.B. provided computational insights. L.M. supervised the project. All authors have read and approved the final version of the manuscript.

