## Supplementary figures and tables for "BI-SCUDO Regression - A novel algorithm to infer clinical variables from omics datasets"

November 17, 2026

| **Parameter** | **Value / Configuration** | **Description / Search Bounds** |
|---|---|---|
| Filtering Threshold ($p$) | Continuous variable | Bounds: $[0, 0.05]$ (Wilcoxon $p$-value) |
| Up-regulated Signature Length ($L_{\text{up}}$) | Integer variable | Bounds: $[10, 200]$ |
| Down-regulated Signature Length ($L_{\text{down}}$) | Integer variable | Bounds: $[10, 200]$ |
| Percentile Normalization Level | Discrete choice | Candidates: $\{0.10, 0.15, 0.20, 0.25, 0.30\}$ |
| Number of Nearest Neighbors ($k$) | Integer variable | Bounds: $[3, 15]$ |
| Population Size | 5,000 | Number of individuals per generation |
| Maximum Generations | 150 | Termination limit |
| Stall Generations | 20 | Early stopping trigger on fitness plateau |
| Elite Count | 20 | Top individuals preserved unchanged (0.4%) |
| Crossover Fraction | 0.8 | Probability of recombination ($P_c$) |
| Mutation Probability | 0.2 | Probability of mutation ($P_m$) |
| Crossover Operators | Blend / Swap | Blend (threshold), Integer Swap ($L_{\text{up}}, L_{\text{down}}$), Uniform ($k$, percentile) |
| Mutation Operators | Gaussian / Discrete | Adaptive Gaussian (threshold), Discrete Random Walk ($n_1$, $n_2$, $k$) |

Table **S1**: Configuration parameters and search space bounds for the Genetic Algorithm (GA) hyperparameter optimization framework in BI-SCUDO Regression.

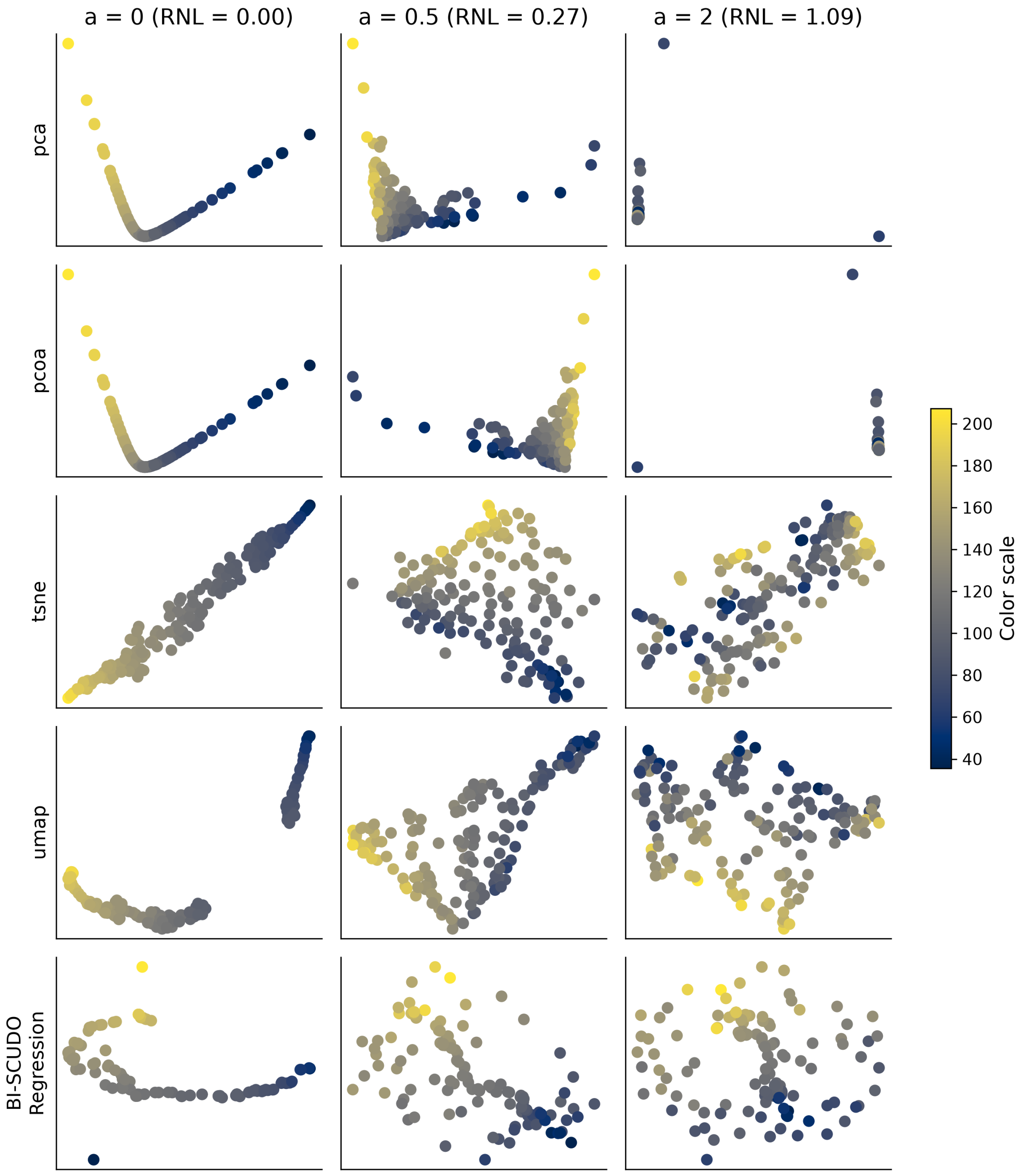


Figure **S1**: **Impact of Noise on the quality of the dimensionality reduction** Comparison of dimensionality reduction techniques (PCA, PCoA, t-SNE, UMAP, and BI-SCUDO Regression) applied to the same dataset under increasing noise levels ($a = 0, 0.5, 2$). Each point represents a sample in a 2D projection colored according to the clinical variable to be predicted. PCoA and PCA offer the best projection at low noise level, but their sensitivity to outliers makes them not suited for noisy dataset. t-SNE and UMAP capture the gradient of the clinical variable at low noise levels, but this structure progressively degrades as noise increases, while BI-SCUDO Regression, even though it is less clear when noise is low, retains some of the underlying structure even when noise becomes prevalent.
